# Low-cost DIY bioreactor system for educational and research applications: an accessible off-the-shelf bubble column concept

**DOI:** 10.1101/2025.04.21.649716

**Authors:** Antoni Gandia, Steven Christiansen

## Abstract

The development of efficient, cost-effective bioreactor systems is pivotal in biotechnological processes involving the production of microbial products like pharmaceuticals, bio-fuels, and protein-rich biomass. Many bioreactor models are commercially available in all industries; however, prices are usually exorbitant, making them inaccessible to low-budget research, schools, and start-up companies. This article introduces a low-cost, DIY modular LSF bioreactor system intended to cultivate filamentous fungi and other microorganisms following a minimalist bubble column approach. Using standard sanitary tri-clamp (TC) fittings common in hygienic industries and household brewing setups, this assembly offers an affordable, adaptable, reparable, and robust solution for microbial production at a fraction of the cost of commercial alternatives. In addition, a wide range of budget-friendly sensors and controllers can be integrated to monitor and automate key parameters, making these bioreactors equally accessible and functional. This bubble column bioreactor concept is unique in its scalability and customization possibilities, significantly reducing research and prototyping costs.

## 1 Introduction

Bioreactors are essential in biotechnology research and industrial bioprocessing, serving diverse fields, from pharmaceuticals to environmental sciences. They ensure controlled conditions for optimal microbial growth and production[1–7]. However, their widespread adoption has been hampered by the prohibitive cost of commercial bioreactors, often ranging from 10,000 to 40,000 USD per unit in the 2-to 10L working volume range (based on current quotations). This financial barrier is especially challenging for academic institutions, citizen science labs, and start-ups, where resources are often limited. The high cost extends beyond the bioreactor itself, encompassing ancillary equipment such as autoclaves and laminar flow cabinets necessary to maintain sterile and axenic conditions, a prerequisite for handling most microbial cultures in the laboratory.

The recent surge in open source hardware and software advancements, especially in affordable 3D printing technology, electronic prototyping tools, and the emergence of programmable microcontrollers such as Arduino, ESP32, and Raspberry Pi, has catalyzed rapid prototyping and innovation across various scientific disciplines[8–14]. This trend has also extended to the development of low-cost do-it-yourself (DIY) bioreactors[15–20]. Among these, an increasing number of projects focus on morbidostat and turbidostat-type bioreactors, which are continuous culturing devices derived from chemostat technology[21, 22]. Notable examples include the eVOLVER[23, 24], the Omnistat[25], Chi.Bio[26, 27], and the most recent EVolutionary biorEactor (EVE)[28]. The latter has even inspired a commercially available open source system known as the Pioreactor[29]. In addition to continuous culture systems, Pechlivani et al.[20] introduced a miniaturized 3D printed stirred-tank bioreactor that integrates Internet of Things (IoT) functionalities and is compatible with standard 12-well plates, allowing real-time monitoring via a mobile app. This system builds on previous designs such as SpinΩ[30] and Spinfinity (Spin ∞)[31], offering a low-cost alternative for small-scale microbial cultivation.

Beyond these examples, a variety of other open source and 3D printed bioreactor designs have emerged, each exploring different engineering approaches to microbial cultivation. For example, Cohen et al.[32, 33] developed the Innocell bioreactor, which utilizes a rotating disk system to cultivate Kombucha bacterial cellulose films. Gabetti et al.[34] introduced a 3D printed perfusion bioreactor, while Schneider et al.[35] presented a miniaturized 90 mL stirred tank bioreactor equipped with integrated single-use pH, dissolved oxygen (DO), and biomass sensors. In a comparative study, Supramani et al.[36] examined an Air-L-shaped bioreactor (ALSB), demonstrating its cost-effectiveness relative to conventional stirred tank reactors (STRs). More recently, Achleitner et al.[37] introduced an autoclavable, 3D printed polylactic acid (PLA) vessel compatible with commercial bioreactor systems. Earlier contributions to cost-effective bioreactor design include the work of Sathiyamoorthy & Shanmugasundaram in 1994[38] and Gueguim Kana et al. in 2003[39], who developed low-cost bioreactors that successfully supported bacterial growth. These innovations highlight the increasing potential of budget-friendly bioreactor designs for a broad range of microbial cultivation applications.

Drawing inspiration from these previous DIY bioreactor developments and in response to the financial bottleneck of acquiring commercial bioreactor system of larger volumes, we would like to introduce a minimalist bubble column (BC) bioreactor assembly that places a premium on modularity and adaptability, allowing users to customize it to meet the specific needs of their microbial cultivation projects without the pre-requirement of owning a 3D printer or having to develop complex skills. The modular bioreactor system presented here is suitable for the cultivation of filamentous fungi, and moreover, the versatility of the assembly theoretically extends to various other microorganisms, including algae, bacteria, yeast, or even combinations thereof[40, 41]. The bioreactor follows a BC configuration to perform aerobic liquid-state fermentation (LSF), which constitutes a well-known approach for the submerged cultivation of microbial cultures[42–45]. Unlike traditional bioreactor systems, which are based on proprietary technology, the concept introduced here leverages a simple assembly of readily available off-the-shelf sanitary tri-clamp (TC) fittings, also known as tri-clover. These fittings follow the ISO 2852, DIN 32,676 and BS 4825-3 standards, and are normally used in hygienic industries, such as brewing and food production[46, 47]. The use of these TC fittings results in an affordable, adaptable, reparable, and robust bioreactor system that dramatically reduces the economic barrier to microbial production, offering a sustainable and accessible alternative to its expensive commercial counterparts.

Throughout this article, we also demonstrate the effectiveness of our DIY BC bioreactor assembly by successfully cultivating industrially and culturally important fungi, including *Aspergillus oryzae* (aka Koji mold)[48], *Rhizopus oligosporus* (Tempeh mold)[49], *Ganoderma sessile* (Trojan conk)[50] and *Psilocybe cubensis* (Magic mushroom)[51–53]. It consistently maintains stable environmental conditions and achieves biomass yields that are on par with those obtained using significantly more expensive alternatives. Furthermore, this article aims to provide a comprehensive overview of how to assembly and use the BC bioreactor in two different volumes, and discusses its potential applications in research and education. Although we acknowledge that the concept introduced here may lack most of the advanced features of commercial systems, such as pH control and sensor-based data logging, it offers unique opportunities for hands-on learning, experimentation, and further improvement. With its simplicity, user-friendly function, and affordability, this system has the potential to democratize the way we approach bioreactor-based microbial cultivation and bioproduction in both academic and professional settings.

## 2 Materials and methods

### Bioreactor components, assembly, and working principle

The basic assembly of a BC bioreactor unit, with recommended volumes between 1 to 5 liters (L), can be built for less than 400 € (see Table 1). For the purpose of demonstrating the functionality of the concept, we assembled two units with working volumes (WV) of 2 and 2.5 L, each having a different height-to-diameter (H/D) ratio. The 2 L version, which has a low H/D ratio of 2.3, is shown in Figures 1a and 1b. For this 2 L built, we used 4-inch TC top and bottom plates, along with a 23 cm long sight glass and suitable 1.5-inch barbed top ports and aeration stone fittings, according to the schematics in Figure 1c. In contrast, the 2.5 L version tested in parallel uses 3-inch TC plates mounted on a 50 cm long sight glass, having a higher H/D ratio of 6.25. It is important to note that increasing the H/D ratio can enhance the gas holdup and mixing efficiency up to a certain point, beyond which further increases may lead to diminishing returns or operational challenges. Therefore, selecting an appropriate H/D ratio should be based on the specific process requirements and performance objectives of the bioreactor.

**Table 1.**
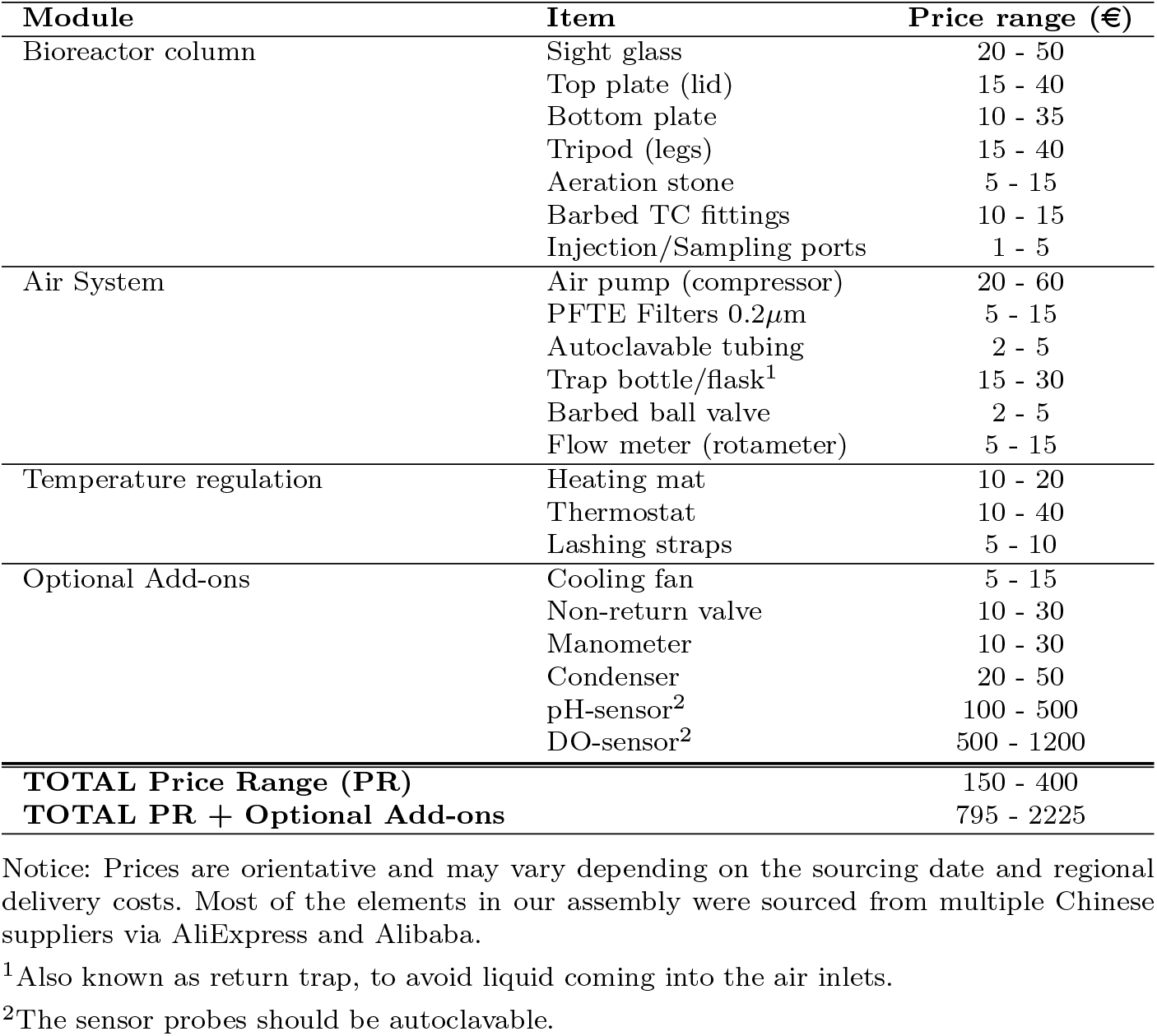
Price list for a basic built.

**Fig. 1.**
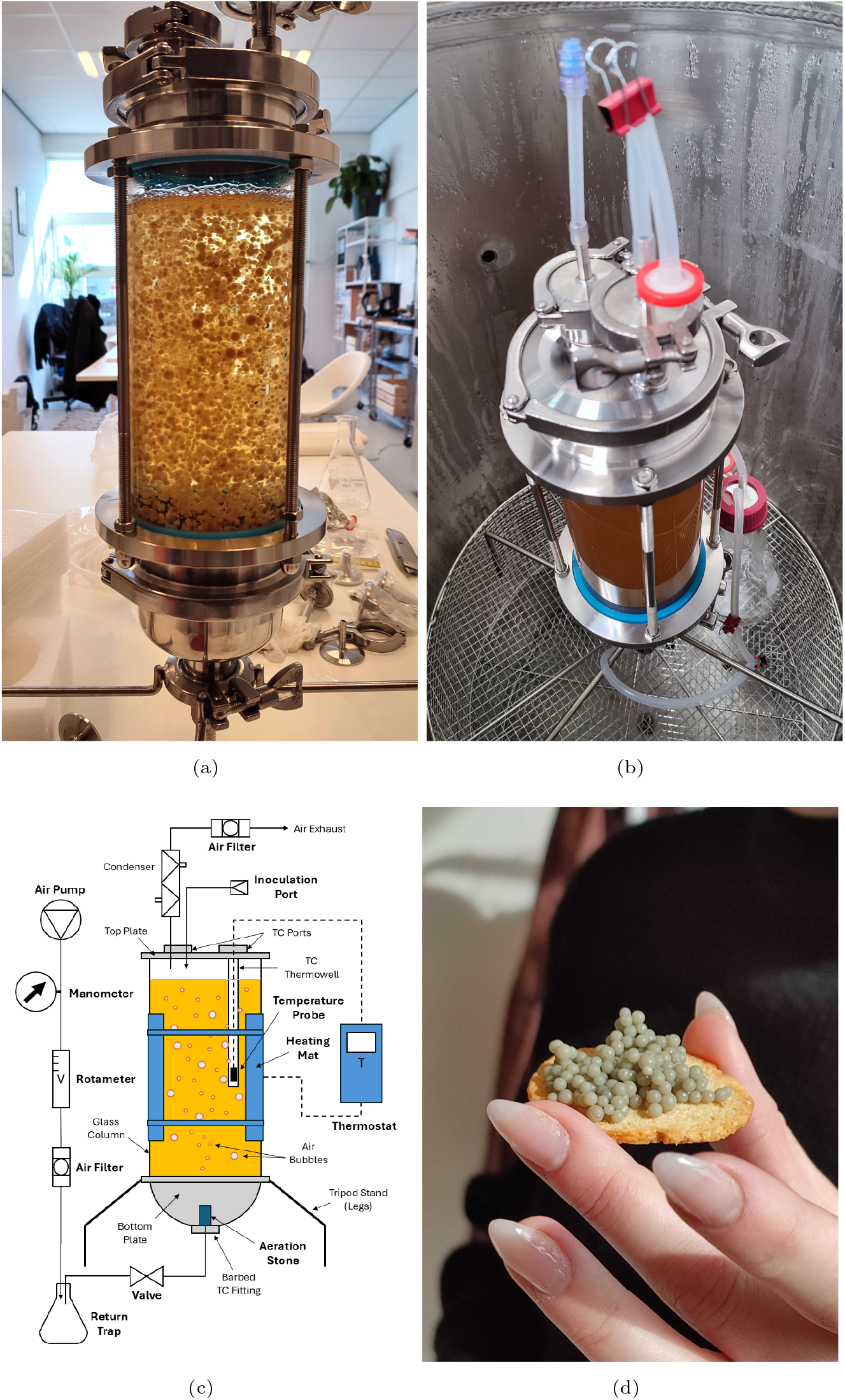
Low-cost bubble column (BC) bioreactor system built using sanitary TC fittings; (a) 2 L BC bioreactor assembly during operation, (b) 2 L BC bioreactor sitting in a 60 L autoclave, (c) working principle schematics of the BC bioreactor, (d) detail of the harvested fungal pellets of the species *P. cubensis*.

The wide catalog of sanitary TC fittings currently available in the global market allows multiple bioreactor configurations able to achieve efficient oxygenation, heating, inoculation, and sterilization procedures, with the potential for further customization and improvements as needed by bioprocesses that require fast prototyping and iterativity. This flexibility in configuration and modification allows for the addition of magnetic and mechanical stirring devices[54], automatic feeding systems, or artificial illumination, which is usually needed by photosynthetic microbes or photoregulated bioprocesses. Furthermore, the system flexibility would allow for the integration of budget-friendly sensors and controllers such as pH sensors, optical density (OD) sensors[55] and fluorometers[17, 27], allowing real-time monitoring and automation of crucial processes for a small fraction of the price compared to their commercial counterparts.

The bubble column reactor configuration (BC) was chosen because it is known for its simplicity in design and operation. It consists of a vertically oriented vessel filled with a liquid phase, with a gas distributor at the bottom, also known as the sparger. This simple structure, which devoids mechanical moving parts, contributes to their ease of maintenance and operation. One of the key advantages of BC reactors is their exceptional rate of heat and mass transfer[56, 57]. The continuous circulation of gas bubbles within the liquid phase promotes efficient mixing and oxygenation, facilitating the growth of a plethora of aerobic microbes, including yeasts, bacteria, and algae[42, 45, 58–61]. Bubble column reactors are particularly interesting for low-budget setups due to their compact design, which not only saves space, but also reduces capital, sanitation, and overall maintenance costs. Their scalability enables them to accommodate varying production volumes, further enhancing their cost-effectiveness. Our concept brings this flexibility over the board by providing a fully built setup with universally compatible sanitary TC fittings, which ease repairs and customization. We also believe that our system could inspire the adaptation and conversion of regular TC jacketed fermentors to fully operational bioreactors capable of controlling working temperatures and sanitation with improved efficiency.

### Maintenance of fungal cultures

Cultures were maintained in 90 mm diameter vented Petri dishes sealed with Parafilm™ by incubation in the dark at 28 °C on a defined malt extract agar (MEA), containing 2 wt% malt extract, 2 wt% agar, and 0.2 wt% yeast extract and transferred to a fresh plate weekly. The experiments were carried out mainly with two fungal species of the Basidiomycota division, *Ganoderma sessile* and *Psilocybe cubensis*. In secondary experiments, other fungal species such as the Ascomycete *Aspergillus oryzae*, and the Zygomycete *Rhizopus oligosporus* were also included (see Table 2 and Fig. 3 for detailed information).

**Table 2.** Fungal species used in this study, including collection code and supplier.

| Taxonomic Identification & Strain | Phylum | Coll. Code | Supplier |
| --- | --- | --- | --- |
| *Aspergillus oryzae* "W-20" | Ascomycota | W-20 | Higuchi, Japan |
| *Rhizopus oligosporus* "TC" | Zygomycota | 60-18 | MOGU S.r.l., Italy |
| *Psilocybe cubensis* "APE Revert" | Basidiomycota | 92-21A | Mimosa Therapeutics, NL* |
| *Ganoderma sessile* "Trojan Conk" | Basidiomycota | 95-19 | MOGU S.r.l., Italy |

\*This company ceased operations in 2022.

### Growth medium and sterilization

The composition of the liquid growth medium was 2% malt extract, 2% glucose, 0.5% yeast extract, and 0.2% vegetal peptone, dissolved in distilled water, with a pH adjusted to 5.5 with HCl and NaOH. The medium was loaded into the assembled bioreactor cylinder using a funnel and then fully sterilized for 20 minutes at 121 °C and 1.5 psi. The autoclaved bioreactor was cooled to room temperature prior to inoculation with an aliquot of a homogenized liquid culture. The bioreactor system was mounted on a flat surface, such as a laboratory bench or table, and the air pump was connected to the aeration stone. The airflow rate was adjusted according to the oxygen requirements of the microbial strain. The temperature of the growth medium was monitored and controlled by using a thermostat (Inkbird ITC-308, China) and an electric heating mat. The cultivation cycle in the bioreactor ran for 8 days.

### Cultivation in shaken flasks for comparison

Conical culture flasks of 2 L (nominal volume) were filled with 1.2 L of growth medium (working volume) and autoclaved for 20 minutes under the same conditions. The flasks were cooled to room temperature, inoculated, and incubated at 28 °C using an orbital shaker incubator set at 180 rpm. For the basidiomycetes, the growth period was 8 days, while for the Ascomycete and Zygomycete fungi it was 4 days. The composition of the growth medium and the inoculum was the same as that used for the bioreactors.

### Inoculation and operation

The bioreactors were inoculated with 10 g/L of fresh homogenized biomass from 48 h flask cultures. The homogenization step was performed with an autoclavable laboratory blender (Waring, USA). Cultures in the bioreactor were kept at 28 °C with an aeration flow rate of 0.5 to 1.0 vvm for 8 or 4 days depending on the species. The aeration was provided by compressed air at 10 psi. The volumetric flow rate of the air was controlled by a rotameter before being filtered through a 0.2 *µ*m PFTE air filter and then delivered to the vessel through a steel porous diffuser, commonly known as an aeration stone (Fig. 2d). The pH of the medium was not adjusted during the duration of the batch cycle.

**Fig. 2.**
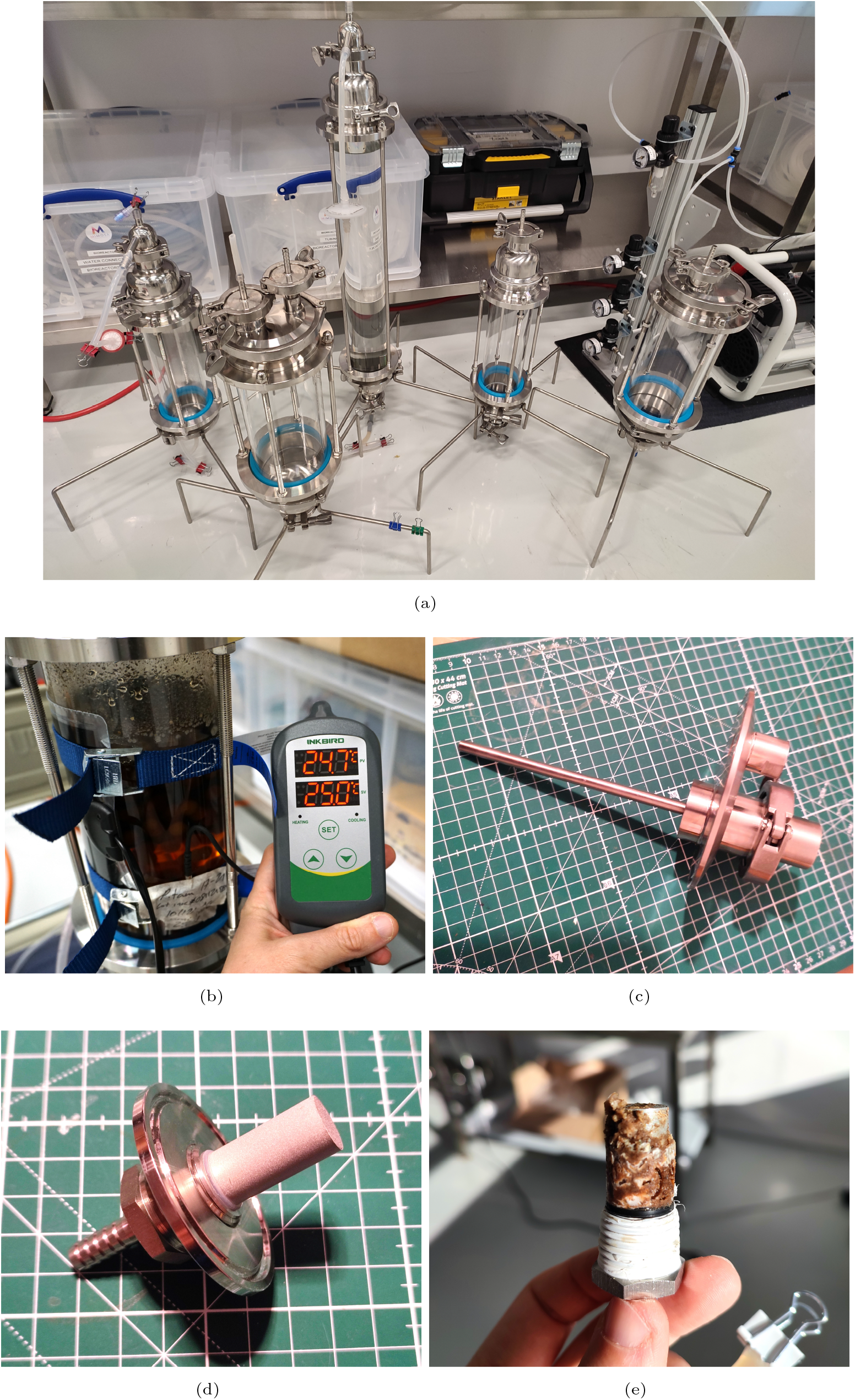
Setup details and versatility: (a) examples of bioreactor prototypes with different volumes, (b) heating mat and thermostat for temperature control, (c) TC thermowell fitting, (d) detail of the aeration stone used in the bioreactor assembly, and (e) aeration stone covered in fungal biomass after a batch.

**Fig. 3.**
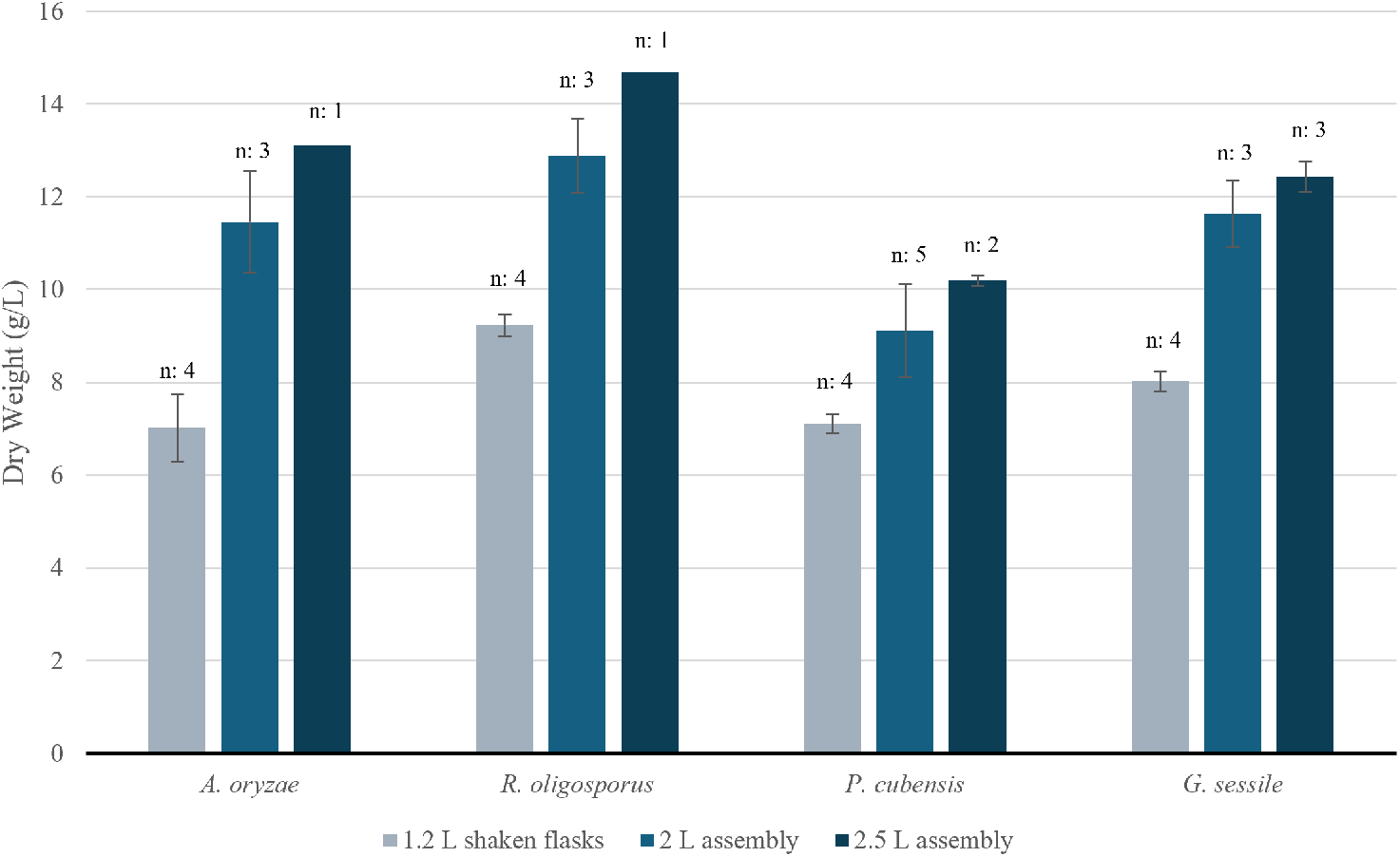
Comparison of yields (DW) between two custom-built BC bioreactor volumes and classical shaken flasks. The number of repetitions is listed on top of each bar (n:).

### Biomass yield estimation

After the cultivation period, which was 8 days for the basidiomycetes and 4 days for the mold species, the wet fungal pellets were filtered using preweighed Whatman grade 1 filter paper and a Büchner funnel connected to a sidearm flask using a neoprene adapter, with a tube leading to a vacuum pump. The collected biomass was rinsed twice with distilled water and then dried at 50 °C until a stable dry weight was achieved. The dry weight (DW) of the biomass was determined by gravimetrically measuring the mass using a digital scale.

### Educational workshop

In order to assess ease of use and educational potential of the low-cost bioreactor, a hands-on workshop was organized at K-Lab Zurich, a community laboratory dedicated to education and practice in mycology and mycelium-based technologies. The workshop, attended by 12 participants with diverse backgrounds in biotechnology and engineering, provided practical training in the assembly, sterilization, inoculation, and operation of a batch process of bioreactors. Participants were first introduced to the modular design of the bioreactor and the function of its components through a guided assembly session. This process emphasized the importance of each element in maintaining operational efficiency and reliability.

In the second phase, participants prepared the bioreactor for fungi cultivation, which included sterilization of the medium-laden vessel using an autoclave and inoculation with a *R. oligosporus* liquid culture. The final phase involved the introduction of a batch fermentation process, in which participants monitored operational parameters such as temperature and aeration to understand their influence on microbial growth. The workshop emphasized the practical application of sterile techniques and bioprocess engineering principles while encouraging active engagement through direct observation and control of the vessels. Participant feedback was collected to assess the educational impact of the workshop and identify areas for improvement, which are reflected in Table 3 and discussed in the results section.

**Table 3.** Pros and cons of the proposed BC bioreactor assembly.

| Advantages | Disadvantages |
| --- | --- |
| Built with standard sanitary TC fittings; no proprietary tech involved, multiple volumes and customization options available. | High foam formation at high gas flow rates. |
| Simple structure; no mechanical stirring required. | Inoculum size and homogenization method can influence consistency. |
| Easier maintenance; no mobile parts and reduced risk of contamination. | Inefficient mixing with high cell densities: growth of pellets in volume and weight over extended periods of time would require continuous aeration adjustments. |
| Low energy requirement; no mechanical agitators or complex electronics. | Large fungal pellets may complicate sampling procedures and reduce yields. |
| Low shear stress; gentle on sensitive cultures. | Presence of "dead zones" that trap biomass; fungal growth can obstruct aeration stones, reducing oxygenation. |
| Comparable yields to commercial bioreactors cited in literature. | Requires a large autoclave for sterilization. |
| Higher yields than shaken flask cultures. | Addition of extra sensor arrays for pH, DO, and optical density, could significantly increase the price tag of the assembly. |
| Sturdy construction; can be enhanced with protective fittings and jacketed vessels. | Lack of advanced computerized controls at this stage of development. |
| Stainless steel fittings; superior to 3D printed tools for durability, withstands repeated autoclaving cycles. |  |

## 3 Results

### Evaluation of temperature control capabilities

The bioreactor temperature control system was evaluated in four different batches using a heat mat coupled with an external thermostat system (Fig. 2b). The heat mat installed on the 2 L vessel was 15×28 cm in size and consumed 7 watts, on the other hand, the heat mat installed on the 2.5 L vessel was 42×28 cm and consumed 20W. The mats were tightly attached to the sight glass with lashing straps. The temperature probe of the thermostat device probe was inserted into a thermowell located on the top plate of the bioreactor, which was filled with glycerol, ensuring precise and safe temperature measurements (Fig. 2c). The thermostat system effectively regulated the heat output of the mat, maintaining a stable internal temperature within the bioreactor. This configuration allowed for accurate temperature modulation, demonstrating the system’s capability to create controlled thermal conditions suitable for microbial cultivation. The combination of the heat mat and the thermowell setup provided consistent thermal stability, highlighting its suitability for small-scale fermentation studies and other temperature-sensitive processes.

### Growth of fungal biomass

The budget-friendly BC bioreactor system demonstrated effective support for the growth and cultivation of four fungal species. The biomass yield was measured in terms of dry weight (DW) after 8 days of cultivation, with values comparable to those obtained in more advanced bioreactor systems[60, 62, 63]. Specifically, the yields of *A. oryzae, R. oligosporus, P. cubensis* and *G. sessile* in the BC system ranged from 8.02 to 14.7 g/L for the 2 L and 2.5 L assemblies, clearly outperforming classical 1.2 L shaken flasks, where the yield ranged from 6.23 to 9.56 g/L. In particular, the yields were slightly higher for the 2.5 L BC assembly, which may indicate a more efficient H/D ratio for fungal cultivation. Furthermore, we confirmed that the heating and aeration provided by the BC assembly was sufficient to maintain stable growth conditions, as reflected in consistent biomass yields in multiple tests.

### Educational workshop outcomes

The workshop results highlighted the low-cost bioreactor as an accessible and effective educational tool. Participants reported a substantial improvement in their understanding of bioprocess engineering, particularly regarding the modularity and transparency of the bioreactor, which enabled direct observation of fungal growth and agitation through the bubble column principle (BC). Post-workshop surveys revealed that 10 of 12 participants found the assembly process straightforward and engaging; on the other hand, 6 participants remarked the challenges and bottlenecks in the sterilization step, which requires a considerable volume of autoclaves, indicating the need for additional improvements and simplification in this regard.

The inoculation and batch fermentation segments were particularly well received, as they allowed participants to understand and handle fungi in a practical way, providing insight into the relationship between operational parameters and microbial growth. Participants also noted the bioreactor’s potential for use in both research and educational contexts, with discussions highlighting its scalability for larger processes and adaptability to various microbial systems other than fungi. In general, the workshop effectively demonstrated the value of the bioreactor in biotechnology education, equipping participants with practical skills and theoretical knowledge essential for microbial cultivation and control of bioprocesses.

## 4 Discussion and Future Prospects

The development of a modular and budget-friendly bioreactor system as detailed in this study marks a significant step towards democratizing access to liquid-state microbial cultivation for educational and research purposes. Using open-source technology principles and building upon the foundation of previous low-cost bioreactor innovations, our concept not only offers an affordable solution but also embodies versatility and adaptability. Our bubble column bioreactor allows the cultivation of various species of filamentous fungi and potentially other microbial species such as yeasts, bacteria, and algae, underlining its utility in various scientific domains. The assembly of the system, which emphasizes simplicity and the use of off-the-shelf components, positions it as a practical tool to foster hands-on learning and experimental research, even in settings with limited resources.

However, the journey toward refining and optimizing this bioreactor is ongoing. Although the system has proven effective for basic cultivation needs, such as the production of fungal biomass in the form of spherical pellets and the achievement of yields comparable to more advanced systems reported in the literature, it may face limitations when dealing with more recalcitrant microbial strains that require advanced sensor and control arrangements. Proper sterilization of the vessel is also another constraint, currently requiring the use of a large volume autoclave able to fit in the entire bioreactor (Fig. 1b), which represents a costly ancillary piece of equipment not reflected in our budget. Alternatives to the use of autoclaves should be investigated, such as, for example, the implementation of mechanisms for on-site sterilization using TC heating elements, chemical sanitation, gamma irradiation, or plasma sterilization methods[64], in combination with filter sterilization of the liquid media. Reducing the overgrowth on bioreactor internal walls, joints, and sensors, testing the influence of different height-to-diameter ratios, and improving the rheological dynamics, such as oxygen transfer and mixing time, should also be taken into account. In addition, installing non-return valves on the air supply lines, which act as barriers against liquid backflow into the air supply system, would reduce such critical risk when operating larger liquid volumes. These challenges underscore the necessity for continuous improvement and adaptation of the system, integrating advanced features and controls that could broaden its applicability.

Furthermore, the possibility of integrating open-source hardware and software, as evidenced by the successful utilization of microcontrollers and IoT functionalities for real-time monitoring in previous open-source bioreactor projects, sets a promising direction for improving the capabilities of the system and addressing its current limitations. In this regard, the addition of infrared (IR) optical density sensors to record biomass growth in real time[65], LED light arrays to grow photosynthetic microorganisms[66], or built-in optogenetic systems[67], could expand the ability of the bioreactor assembly to serve scientific research and analytics properly. Furthermore, we recommend studying the effects of the height-to-diameter ratio and the aeration rate on dissolved oxygen levels and the dynamics of pellet formation[68]. We also recommend further research on the optimal conditions for cultivating different species, such as adequate carbon and nitrogen sources and ratios, inoculum density, temperature, and pH.

Looking ahead, the potential of this simple bioreactor system extends beyond its current configuration and applications. The prospect of further research and development to optimize its performance for a wider array of microbial products is compelling. Such endeavors could democratize processes ranging from single-cell protein production to waste treatment and biochemical synthesis, making advanced biotechnological applications more accessible to a broader audience. The integration of the IoT-based controls mentioned above and the exploration of up-scaling options, such as, for example, the refurbishment of larger jacketed brewing-type fermentors, underscore the potential of the system for evolution, aiming not only to meet but to anticipate the future needs of both academic and industrial biotechnology sectors. Finally, we remark that this initiative embodies a commitment to innovation, accessibility, and the advancement of science through community-driven efforts, expecting feedback, assembly iterations, and overall improvements from the broader DIY biosciences community, which should collectively pave the way for a more inclusive and collaborative approach to bioreactor technology.

## Declarations

### Data availability statement

The datasets generated during and/or analyzed during the current study are available from the corresponding author upon reasonable request.

### Author contributions

A.G. conceived and designed the bioreactor. A.G. collected the data and wrote the original draft. S.C. supervised, reviewed, and edited the draft. Both authors have read and agreed to the final version of the manuscript.

### Compliance with regulations

Experiments involving the cultivation of *Psilocybe cubensis* were conducted in the Netherlands following the current legal regulations of the country on the handling and study of psychoactive fungi.

### Funding

This work was supported by Experiment Foundation, Footprint Coalition, Schmidt Futures, and the Gordon and Betty Moore Foundation. Grant: Low-Cost Tools for Science (2023).

## Acknowledgments

The authors would like to thank the following people for their endorsement and donations to build the bioreactor prototypes; Stephen Apkon, Maurizio Montalti, Alessandro Chiolerio, Mohammad Mahdi Dehshibi, Jara Salueña Martin, Craig Ellins, Elena Baltaga, Alice Nušlová, Justin Fornelius, Thibaud Trichereau, Aart Van Bezooijen, Gerrit Niezen, Miles Adams, Martin Currie, Olaf Zumpe, Dirk Ehrenberg, Maya Benami, James Keim, Guillaume Costa, Paige Perillat-Piratoine, Maria Higelin, and Rita Hamm. Special thanks to Sebastian Dörner, Felix Blei, Amanda Feilding, and Bob Otis Stanley for their invaluable contributions during the conceptualization and testing phase. Additionally, we are grateful to Andrew Adamatzky from the Unconventional Computing Lab (UWE Bristol, UK) for critically proofreading the manuscript. We also appreciate MOGU Srl, Mimosa Therapeutics PBC, and Higuchi Matsunosuke Shoten Co.,Ltd. for providing the strains used in our experiments. Lastly, we thank Linda Niewold for her assistance in capturing the close-up image of fungal pellets featured in this manuscript.

## Conflict of interest

The authors declare that they have no identifiable competing financial interests or personal relationships that could reasonably be construed as having an impact on the research described in this article.

## Notes

### Competing Interest Statement

The authors have declared no competing interest.

